# Single-domain antibodies targeting the pan-T-cell markers CD2 and CD7 as universal immunotherapy T-cell tracers

**DOI:** 10.64898/2025.12.23.693936

**Authors:** Dario Gosmann, Lisa Russelli, Theresa Käsbauer, Sandro Bissenberger, Florian Bassermann, Wolfgang A. Weber, Calogero D’Alessandria, Angela M. Krackhardt

## Abstract

**Rationale:** Despite significant advancements in immunotherapeutic approaches, such as immune checkpoint modulation and CAR-T cell therapy, reliable and universally applicable surrogate markers to monitor and evaluate the wide variety of therapeutic responses are still missing. This gap of information can lead to delayed assessment of responders versus non-responders and to misinterpretation, potentially resulting in premature or erroneous treatment decisions. Thus, a universal surrogate marker may be of preeminent value in monitoring immune responses, guiding immunotherapies, and ultimately improving patient outcomes. To address this challenge, we developed two immunoPET tracers based on single-domain antibodies (sdAbs) that target the pan T-cell markers CD2 and CD7 to monitor and visualize T cells distribution in vivo.

**Methods:** This study aimed to establish a comprehensive approach for clinical translation by assessing the feasibility and key characteristics of CD2- and CD7-sdAb for safe, effective clinical application. Binding specificity was confirmed through antigen transduction and knockout in tumor and T-cell leukemia cell lines. In vitro, cytokine secretion and T-cell cytotoxicity were evaluated, while a xenogeneic myeloid sarcoma mouse model was used to analyze T-cell cytotoxicity and visualize TCR-transduced CD8^+^ T cells at the tumor site using PET/MR imaging.

**Results:** This study demonstrates the high specificity, affinity, and thermal stability of CD2- and CD7-sdAb, enabling efficient radiolabeling without impairing T-cell functionality. Specifically, binding to human CD8⁺ T cells did not alter cytokine secretion or cytotoxicity in vitro. Furthermore, in vivo, the cytotoxic ability of transferred TCR-transduced human CD8^+^ T cells was not compromised upon i.v. application of CD2- and CD7-sdAb.

Ultimately, intravenously injected ^68^Ga-NOTA-CD2-sdAb and ^68^Ga-NOTA-CD7-sdAb were respectively able to clearly depict and visualize previously administered TCR-transduced CD8^+^ T cells at the tumor site by PET/MRI.

**Conclusions:** In this study we have developed two pan T-cell tracers that are able to visualize human T cells and clearly differentiate between responding- and non-responding tumors while not impacting T-cell functionality in vitro and in vivo. The results provide a base for clinical use of these tracers as pan T-cell tracers applicable for any form of immunotherapy, making them a crucial tool in guiding immunotherapeutic decision-making.

## Introduction

The revolutionary development of cancer immunotherapies such as immune checkpoint inhibitors and adoptive transfer of T cells, as e.g. CAR-T, has changed the way various solid and hematologic malignancies can be treated. Particularly immune checkpoint blockage (ICB) targeting PD-1 / PD-L1 [1] and CTLA-4 [2] as well as anti-CD19 CAR-T cells [3] but also the application of multi-functional tumor-targeting T-cell recruiting antibodies [4] have seen resounding success over the past years. However, immunotherapies are highly diverse in terms of their target structures and mechanism of action. Moreover, response and progression patterns differ widely from conventional anticancer treatments [5] and therefore evaluation of responses is critical. Hard to assess immunotherapeutic responses such as pseudoprogression [6, 7], hyperprogression [7, 8] as well as delay of progression and effects on subsequent treatment [9] have been described in various tumor entities, especially in melanoma [10], but also for HNSCC [11], renal-cell carcinoma [12] and NSCLC [13–15].

After more than a decade of immunotherapy success, sensitive surrogate biomarkers that allow for timely evaluation of immunotherapeutic response patterns, their pharmacodynamics and- kinetics as well as treatment termination decision criteria are still missing and necessary to improve outcomes for patients receiving immunotherapy [16]. Current options to monitor and evaluate immune reactivity and response of patients include lymphocyte analysis from whole blood or tumor-biopsies, which each come with significant drawbacks. While analysis of lymphocytes can gather information on genomic, transcriptomic and proteomic status of immune cells [17, 18], monitor cytokine secretion profiles and further markers found in the serum [19–21], it fails to provide real-time information about tumor-specific localization, infiltration and activation of immune cells [22, 23]. Biopsies on the other hand can provide tumor-specific insights, yet, can only partially and locally depict the dynamic and heterogeneous immune responses. Moreover, an invasive procedure is required. Due to these inherent limitations of ex vivo analyses, non-invasive imaging technologies such as ^18^FDG-PET emerged as faster and more expedient approaches to assess immunotherapeutic success. However, even though ^18^FDG-PET offers high sensitivity and functional information as a routine tool in clinical oncology [24], it is limited due to the fact that an increase in metabolic activity may be induced by tumor growth but also infiltrating immune cells, which may result in interpretive errors [25].

To solve the problems that arise from estimating the patients multifaceted immune-responses passively via circulating blood markers or ^18^FDG-PET, directly monitoring the most significant immune cells involved in antitumor-immunity would be ideal. Current research on this subject is mainly focused on imaging individual T-cell subsets such as CD8^+^ T cells [26] rather than the overall T-cell population. While tumor-infiltrating CD8^+^ T cells have long been shown to correlate with improved prognosis in a variety of tumor entities [27–29], these narrowly defined populations fail to depict the versatile aspect of T-cell responses during immunotherapy to its full extent. For instance, various studies have shown the added predictive value of CD4^+^ T cells in the outcome of ICB across multiple entities [30–32] and current research is suggesting a more dominant role for CD4^+^ T cells during immunotherapies [33–35], highlighting the importance of an encompassing observation of the whole T cell population. In addition to ICB therapy, CD4^+^ T cells have been proven to be of high relevance for the success of CAR T cell therapy in approved CAR T cell products like lisacabtagen-maraleucel [36], as well as diverse preclinical studies in glioblastoma [37] and leukemia patients [38], and furthermore CD4^+^ cells may be of particular importance for long-term tumor eradication. Taken together, the significance of encapsulating the whole spectrum of immunotherapeutic responses is fast-growing, and with it the need to monitor and evaluate the immune response during therapy with a universally applicable surrogate marker.

For this purpose, we have developed two independent single-domain antibody (sdAb)-based pan-T-cell tracers to directly depict both, endogenously stimulated as well as genetically modified T cells during the course of immunotherapy. This is made possible by the unique properties of the targeted cell-surface markers, CD2 and CD7, which are not only expressed on all T-cell subsets but whose expression is furthermore upregulated on specifically activated T cells [39]. These features enable our CD2- and CD7-targeting sdAb to visualize T cells irrespective of subsets or genetic modifications and to highlight the key component in immunotherapeutic responses, specifically activated T cells.

## Methods

### Primary material and cell lines

After informed consent following requirements of the local ethical board and principles of the Helsinki Declaration, whole blood from healthy patients was collected for isolation of peripheral blood mononuclear cells (PBMC) by density-gradient centrifugation. Isolation, stimulation and cultivation of cells was performed as previously described [40–42].

CD8^+^ T cells were negatively isolated using the Dynabeads™ Untouched™ Human CD8 T Cells Kit (Thermo Fisher Scientific) following the manufacturer’s recommendations. The use of CD8⁺ T cells for in vitro assays was based on their cytotoxic role in targeted cellular immune response and the ease of establishing functional tests. However, since CD2 and CD7 are expressed on CD8^+^ as well as CD4^+^ T cell subpopulations, the tracers are expected to reliably detect CD4⁺ T cells as well. The following cell lines were used in this study: human acute leukemia cell line ML2 (The CABRI consortium), B-cell lymphoma cell line U-698-M (DSMZ), human acute promyelocytic leukemia cell line NB4 (Cell Lines Service, CLS), acute myeloid leukemia cell line HL60 (CLS), melanoma cell line 624.38 Mel (kindly provided by E. Noessner, Munich, Germany), acute T cell leukemia cell line Jurkat, Clone E6-1 (kindly provided by J. Ruland, Munich, Germany) and retroviral packaging cell line 293Vec-RD114 (BioVec Pharma). Cell lines were cultured as previously described [43] and routinely tested for mycoplasma infection (Venor^®^GeM mycoplasma detection kit, Minerva Biolabs).

ML2, NB4 and HL60 cells were retrovirally transduced as previously described [40] with genes coding for HLA-*B*\*07:02 or HLA-*B*\*15:01 respectively, linked to enhanced green fluorescent protein (GFP). Likewise, U-698-M cells were retrovirally transduced with human CD2- and CD7 genes linked to the near-infrared red fluorescent protein (iRFP). CD8^+^ T cells were retrovirally transduced with the previously described MPO_5_-specific TCR 2.5D6 linked to red fluorescent protein (2.5D6iRFP T_CM_) [42, 43] to specifically target MPO-expressing tumor cell lines.

### Llama Immunization and VHH library construction for CD2 and CD7 sdAb discovery

SdAb against human CD2 and CD7 were generated using established immunization and library construction protocols [44]. A llama was immunized weekly for six weeks at the Vlaams Instituut voor Biotechnologie (VIB) in Brussels with ~100 µg of His6-tagged recombinant extracellular domain (ECD) of human CD7 (Ala26-Pro180) (R&D Systems, 7579-CD-050) or recombinant ECD of human CD2 (Lys25-Asp209) fused to a human IgG1 Fc domain (hCD2-Fc) (G&P Biosciences, FCL2552), both formulated with Gerbu adjuvant P. Peripheral blood lymphocytes were used to prepare cDNA, serving as templates for the amplification of variable domains of heavy-chain-only antibodies. The amplified fragments were cloned into phagemid vectors and transformed into *E. coli* TG1 cells, creating libraries exceeding 10⁸ transformants. After three rounds of panning on solid-phase coated antigens, sdAbs were enriched ~700-fold. For CD7, 63 full-length sdAbs belonging to six CDR3 groups were identified, while 19 unique full-length sdAbs were isolated for CD2. Immunization, VHH library construction and sdAb-enrichment was conducted by the VIB in Brussels.

### Expression and purification of sdAb

Anti-CD2 and –CD7 sdAb DNA fragments were recloned from pMECS- into pHEN6c-vectors to retain a carboxyterminal hexahistidine tail [45] and were subsequently transformed in *E. coli* WK6 as described [46]. Transformed *E. coli* WK6 were grown overnight in 10 ml LB-Medium, supplemented with 100 µg / ml ampicillin and 1 % glucose at 37 °C, after which 1 ml of pre-culture was added to 330 ml Terrific Broth medium (Thermo Fisher Scientific), supplemented with 100 µg / ml ampicillin, 0.1 % glucose and 1 mmol / l MgCl_2_. When optical density (OD_600_) of 0.9 was reached, sdAb expression was induced by addition of 1 mM Isopropyl-β-D-thiogalactopyranosid (IPTG) overnight at 28 °C. When the density of culture reached an OD_600_ of 25, bacterial pellets were resuspended in TES-buffer (0.2 M Tris pH 8.0, 0.5 M NaCl, and 0.5 mM EDTA) and stirred for 1 h at 4 °C. sdAb were extracted from periplasm of *E. coli* by centrifuging culture mixture at 8000 × g for 30 min at 4 °C. sdAb were purified using the Thermo Scientific HisPur Ni-NTA Spin Purification Kit, which utilized Ni-NTA spin columns for immobilized metal affinity chromatography (IMAC). Buffer of purified sdAb was exchanged from 250 mM imidazole elution buffer to PBS.

### sdAb analysis via SDS-PAGE, SE-HPLC and ESI-MS

Purity of sdAb was analyzed by SDS-PAGE gel electrophoresis under non-reducing conditions, running a 10 % NuPAGE Bis-Tris Mini Gel (Thermo Fisher scientific) for 30 min at 200 V with subsequent staining using Coomassie Blue staining solution and destaining overnight.

Size exclusion high performance liquid chromatography (SE-HPLC) was performed with a Yarra™ 3 µm SEC-3000 LC column (Phenomenex) using 0.05 M phosphate buffer and 0.15 M NaCl, pH 7.0 as mobile phase at an isocratic flow rate of 1.0 ml·min-1. UV-VIS profiles of the antibody-derivatives were acquired at 280 nm and radioactive detection was performed via a GABI Star γ-detector (raytest). The chromatographic runs were carried out on a Shimadzu HPLC system and data were analyzed with the Chromeleon 6.8 chromatography data system software. Instant thin layer chromatography (iTLC) was performed on glass microfiber chromatography paper impregnated with silic acid (Agilent Technologies) using 0.1 M sodium citrate pH 5 as mobile phase. The read-out of the chromatography strips was performed using a radio-TLC-scanner (Bioscan, Eckert & Ziegler) and data were analyzed by the Bio-Chrom Lite software.

MS experiments were conducted on an LCQ-FLEET (Thermo Scientific) system equipped with a 3D ion trap and using electro spray ionization (ESI). The instrument is connected to a high-performance liquid chromatography (HPLC) system (Thermo UltiMate 3000, column: Dionex with a Retain PEP, Drop-in, 10×2.1 mm).

### Flow cytometry analysis, K_d_ determination and thermostability analysis

The following antibodies, antibody-derivatives and cell labeling agents were used for flow cytometry: The aTCRmu-F(ab’)_2_ was prepared from supernatant of H57-597 hybridoma cells using Protein A-Sepharose (GE Healthcare) and a F(ab’)_2_ Preparation Kit (Thermo Scientific Pierce™). For flow cytometry, anti-human CD2 (clone RPA-2.10, OKT11, BioLegend), anti-human CD7 (clone M-T701, BD Biosciences), anti-human CD4 (clone SK3, BD Biosciences), anti-human CD8 (clone SK1, BioLegend), anti-human HLA-B7 (clone BB7.1, Novus Biologicals), anti-histidine tag (clone AD1.1.10, Bio-Rad Laboratories) and anti-mouse TCR β Chain (clone H57-597, BD Biosciences) were used. 7-Aminoactinomycine (7-AAD, Sigma-Aldrich) was used to identify dead cells, all samples were analyzed using the flow cytometer LSRII (BD Biosciences) and data was evaluated using FlowJoSoftware10.8.0 (FlowJo, LLC).

To determine the dissociation constant (K_d_), CD8^+^ T cells were stained with varied concentrations of CD2-sdAb and CD7-sdAb and analyzed by flow cytometry. The K_d_ was then determined using nonlinear regression analysis of plotted mean fluorescence intensity values versus used sdAb concentrations.

### Thermal stability of sdAbs

Thermal binding stability of sdAb was assessed by incubating them at 37 °C, 60 °C or 90 °C for 0.5 h, 1 h, 2 h, 4 h and 6 h, respectively, while a sdAb that was kept at 4° C was used as control as described [47]. CD2- and CD7-positive Jurkat E6.1 cells were used as target cells and 0.2×10^6^ cells were washed with FACS buffer and then seeded in a 96-well plate. The respective sdAb was added to the sample at a concentration of 5 nM and the cells were incubated for 60 min at 4 °C. Afterwards, cells were washed three times with FACS buffer and 1 µl of anti-His-Tag antibody was added per well and the samples were again incubated for 60 min at 4 °C. Cells were washed with FACS buffer and analysis of the samples was performed on a LSRII (BD Biosciences) and results were analyzed using FlowJo v7.6.5 software (BD Bioscience).

### In vitro analysis of cytotoxic potential and cytokine production of T cells

TCM and cDMEM were mixed in a 1:1 ratio and 200 µl of the mix were added to a 96-well xCELLigence plate. After 30 min incubation at room temperature, a background measurement of the plate was performed on the xCELLigence RTCA MP instrument. Then, 100 µl of medium was carefully removed from each well and 7.5×10^4^ tumor cells, resuspended in the same TCM and cDMEM mix, were added per well. The plate was then incubated at 37 °C for 24 h in the xCELLigence RTCA MP instrument and the cell index was determined every 30 min. After 24 h, 100 µl of medium was carefully removed and 7.5×10^4^ transduced CD8^+^ T cells as well as sdAbs (c_END_ = 100 nM or 500 nM) were added in 100 µl of TCM and cDMEM mix. Each condition was evaluated in triplicates. After another 24 h incubation at 37 °C with a cell index analysis every 30 min, the coincubation was stopped and the data was analyzed using RTCA software 2.0. The cell index was normalized at the time of T-cell addition. Calculation of percentage cytolysis was done using the following formula: 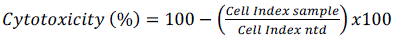.

Cytokine secretion was evaluated after 24 h co-incubation of TCR 2.5D6 CD8^+^ T cells (20,000 cells /well) with MPO-expressing cells derived from ML2-B7, NB4-B7 and HL60-B7 at a ratio of 1:1 in 96-well round bottom plate, with the addition of CD2-sdAb and CD7-sdAb at a concentration of 100 nM and R3b23-sdAb as control. The R3B23-sdAb, which targets the M-protein of 5T2MM myeloma cells derived from C57BL/KaLwRijHsd mice, was provided by VIB and served as a suitable negative control for this study, as it has no expected reactivity with human cells [48]. Supernatants from the co-culture were used to determine secretion levels of IL-2, GM-CSF and TNFa by ELISA, according to manufactureŕs instructions (BD Bioscience).

### In vivo tumor rejection studies

Immunocompromised NSG (NOD.Cg-Prkdc^scid^ Il2rg^tm1Wjl^/SzJ) mice (The Jackson Laboratory) were used for a previously described myeloid sarcoma model [39]. In brief, female NSG mice at age 6 – 12 weeks were subcutaneously injected with 1×10^7^ ML2-B7 and ML2-B15 control tumor cells on the right and left shoulder respectively and after eight days, 2×10^7^ 2.5D6 CD8^+^ T cells were intravenously injected. Three days later, 20 µg of CD2-sdAb and CD7-sdAb were also intravenously injected, with anti-CD2-F(ab’)_2_ (clone OKT11, anti-CD2-F(ab’)_2_ (clone RPA-2.10), R3b23-sdAb and PBS used as controls. Kinetics of tumor growth were monitored daily for 20 days post tumor-cell injection and assessed using a digital caliper. Mice were maintained according to conventional institute guidelines and with the approval of local authorities.

### Conjugation of p-SCN-Bn-NOTA to sdAb

The p-SCN-Bn-NOTA chelator (Macrocyclics) was conjugated to sdAb to allow the complexation of ^68^Ga isotope. To consent the conjugation of the NCS group of the chelator to the -NH_2_ groups of sdAb’s lysins, a buffer exchange was performed using Amicon Ultra-15 centrifuge filter units (3K) (Merck) and 0.05 M sodium carbonate buffer, at pH 8.7. To a 1-2mg/ml solution of sdAb, a 10-fold molar excess p-SCN-Bn-NOTA was added, and the mixture was incubated for 2h at 25°C eventually adjusting the pH with Na_2_CO_3_ 2M. The NOTA-sdAb conjugate was then separated from the unreacted chelator by gel filtration using a pre-rinsed disposable PD-10 column and 0.1M NaOAc buffer solution at pH 5. Quality controls were performed via SDS-PAGE, as previously described, and by SEC HPLC to exclude the presence of aggregates and unreacted NOTA chelator. Size exclusion chromatography was performed on a Superdex 30 increase 3.2/300 column (GE Healthcare) with isocratic method using PBS (pH 7.4) as eluant (NOTA-sdAb R_t_ = 10.4 min).

### Synthesis of ^68^Ga-labeled sdAb

Using a 0.1M HCl solution, [^68^Ga]gallium was eluted from a ^68^Ge/^68^Ga generator (Eckert & Ziegler) as [^68^Ga]GaCl_3_ The most concentrated fraction (1.2ml – 400-600MBq) is used for the radiolabeling reaction.

A solution of 0.6ml of 0.1M HCl containing 200-300MBq of [^68^Ga]gallium was added in a 1M NaOAc pH 5 buffer solution containing 120µg of precursor, NOTA-sdAb previously described, and resulting in a pH 4.5-4.7. The reaction mixture was kept at 25°C for 10min. After this time the product was purified by gel filtration using a disposable pre-rinsed PD-10 column and a solution of NaCl 0.9% + 5mg/ml ascorbic acid as scavenger, at pH 6, freshly prepared.

Quality controls were performed both via instant thin-layer chromatography (iTLC) and HPLC. For iTLC, the stripes were read-out using a radio-TLC-scanner (Bioscan, Eckert & Ziegler) and data were analyzed by the Bio-Chrom Lite software. The HPLC was performed on a system (Shimadzu) with a Photo Diode Array detector (Shimadzu) and a GABI Star γ detector (Raytest). iTLC was performed on silica gel stripes (Agilent) using 0.1 M sodium citrate, pH 5.0, as eluent to evaluate radiochemical purity (RCP) of [^68^Ga]Ga-NOTA-sdAb. Retention factors of the product and unbound gallium-68 are respectively R_f_ = 0 and R_f_ = 1.

Size exclusion chromatography (SEC) was performed on a Superdex 30 increase 3.2/300 column (GE Healthcare) to evaluate RCP (R_t_ = 10.6 min) and absence of aggregates. Isocratic method with PBS (pH 7.4) as eluent.

### PET / MR imaging

All the imaging studies were performed using a nanoScan 3T PET/MRI scanner (Mediso; Budapest, Hungary). Mice were anesthetized using 1.5 % isoflurane and subsequently injected with the tracer into the tail vein. Right before image acquisition, mice were placed on a preheated bed (set at 38 °C) of the scanner device in prone position under constant anesthesia with 1.5 % isoflurane. The initial MRI scan of 10 min was followed by acquisition of static PET emission images for 20 min, after which the mice were sacrificed for further ex vivo analyses. The obtained PET/MRI images were reconstructed, fused and further processed using the Nucline software (Mediso). Tracer uptake was displayed as standardized uptake value (SUV).

### Ex vivo analysis

After PET-MR acquisition or when pre-defined endpoint criteria, animals were sacrificed and tumors as well as organs were removed. The obtained organ samples were employed for subsequent biodistribution analysis. Each tissue sample was weighed and counted using a gamma counter (Wizard 2480, PerkinElmer), together with standards prepared from radiolabeled sdAbs. Accumulation of the tracer was calculated and expressed as the percentage of injected dose per gram of tissue (% ID / g).

### Statistical analysis

Data are presented as mean ± standard deviations (SD). Statistical analysis of results was performed using GraphPad Prism software version V.8.0.2 using a two tailed non-parametric test (Mann-Whitney test) as indicated in Figure legends.

## RESULTS

### Generation and characterization of single-domain antibodies targeting CD2 and CD7

To generate single-domain antibodies (sdAbs) targeting human CD2, a llama was immunized at the VIB with recombinant hCD2-Fc protein. Following immunization, peripheral blood lymphocytes were harvested and used to construct a VHH phage display library (Core 95), comprising approximately 10⁸ independent transformants with a 94% correct insert rate. The library was subjected to three rounds of panning against immobilized hCD2-Fc, resulting in a progressive enrichment of CD2-specific phages: ~15-fold after the first round and up to ~1000-fold by rounds two and three. Of 285 bacterial clones screened by ELISA using periplasmic extracts, 47 clones showed selective binding to hCD2-Fc but not to human IgG1 Fc alone. Sequence analysis identified 19 unique Nanobodies grouped into four CDR3-defined B-cell lineages, reflecting a diverse yet convergent immune response. These CD2-specific sdAbs served as the basis for further characterization and tracer development (Fig. 1A). SDS-Page gel electrophoresis showed a high degree of purity with no visible agglomeration or isoforms for anti-CD2-sdAb (CD2-sdAb) and anti-CD7-sdAb (CD7-sdAb) (Fig. 1B), with the elution profiles (Fig. 1C) for CD2-sdAb (top) and CD7-sdAb (bottom) from the size-exclusion chromatography confirming these results. The precise molecular weights were determined using electro spray ionization based mass spectrometry with preceding high performance liquid chromatography (LCQ Fleet, HPLC-MS) and revealed a molecular mass of 14.180 Da for CD2-sdAb and 14.581 Da for CD7-sdAb (Fig.1D).

**Fig. 1.**
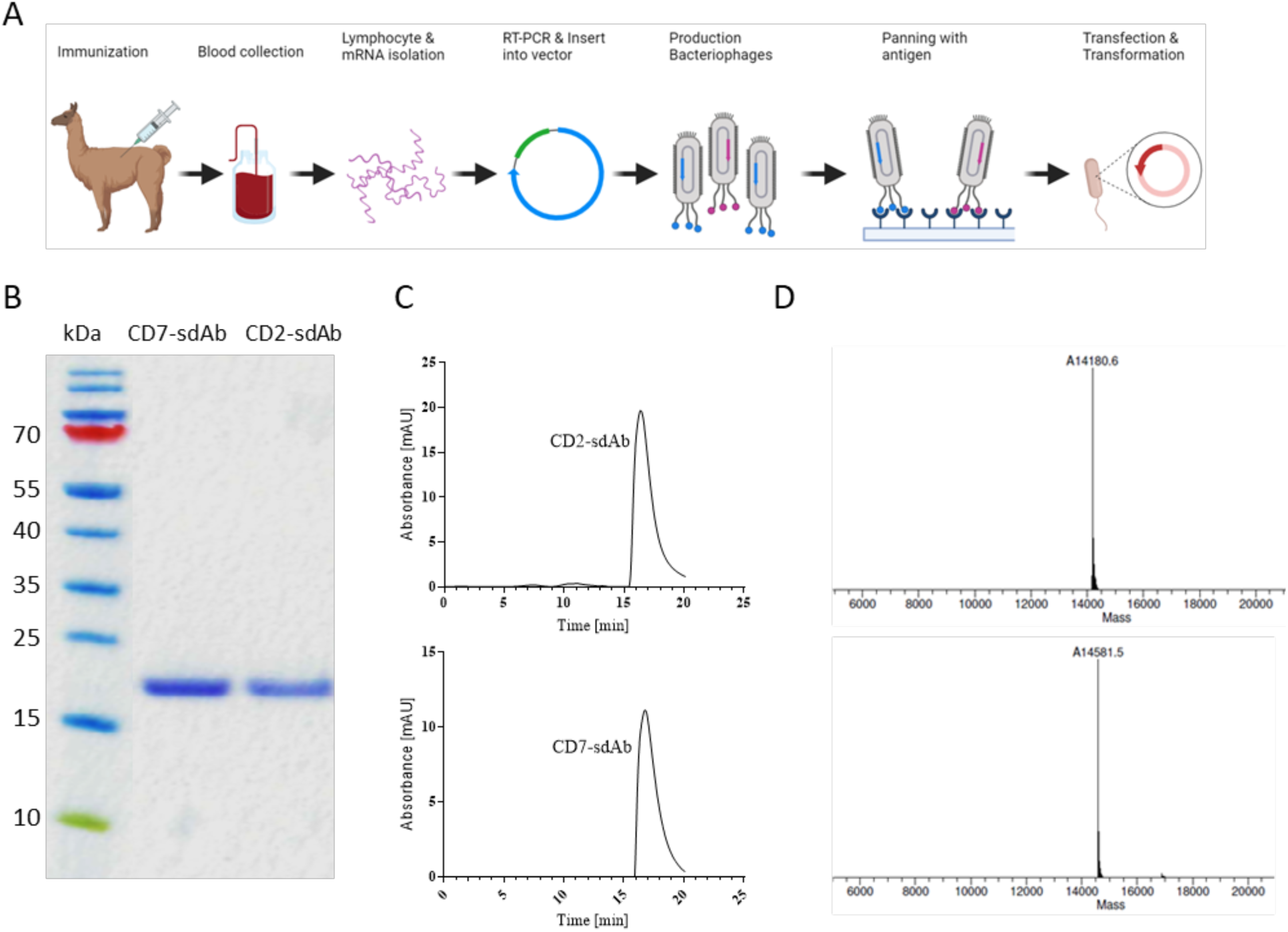
Characterization and validation of CD2-sdAb and CD7-sdAb. **A**, Scheme of generation of sdAb. Llamas are immunized, and lymphocyte mRNA is extracted from collected blood. sdAb sequences are amplified via RT-PCR, cloned into a vector, and used for bacteriophage display. High-affinity binders are selected through antigen panning, followed by bacterial transfection for expression. **B**, SDS-PAGE gel electrophoresis of generated sdAb (L: Size marker, 1: CD2-sdAb, 2: CD7-sdAb). **C**, Size-exclusion chromatograms of CD2-sdAb (top) and CD7-sdAb (bottom). D, HPLC-MS analysis for purity and size determination of CD2-sdAb (top) and CD7-sdAb (bottom).

### In vitro binding studies of anti-CD2 and anti-CD7 sdAb show specificity, high affinity and thermal stability

In order to reliably and selectively depict T cells during a multifaceted immune response amidst the tumor microenvironment, highly specific and stable binding of CD2-sdAb and CD7-sdAb to T cells is essential.

The binding capacity of sdAb to either CD2 or CD7 was assessed twofold. First, specific binding of CD2-sdAb and CD7-sdAb to CD8^+^ human T cells was confirmed by flow cytometry analysis, with a non-targeting irrelevant sdAb (R3b23-sdAb) that served as a reference for non-specific binding (Fig. 2A). Furthermore, low dissociation constants (K_D_) were determined for each construct resulting in K_D_ of 2.30 nM in case of CD2-sdAb and in a K_D_ of 6.34 nM in case of CD7-sdAb, indicating in both cases strong binding to their specific targets (Fig. 2B).

**Fig. 2.**
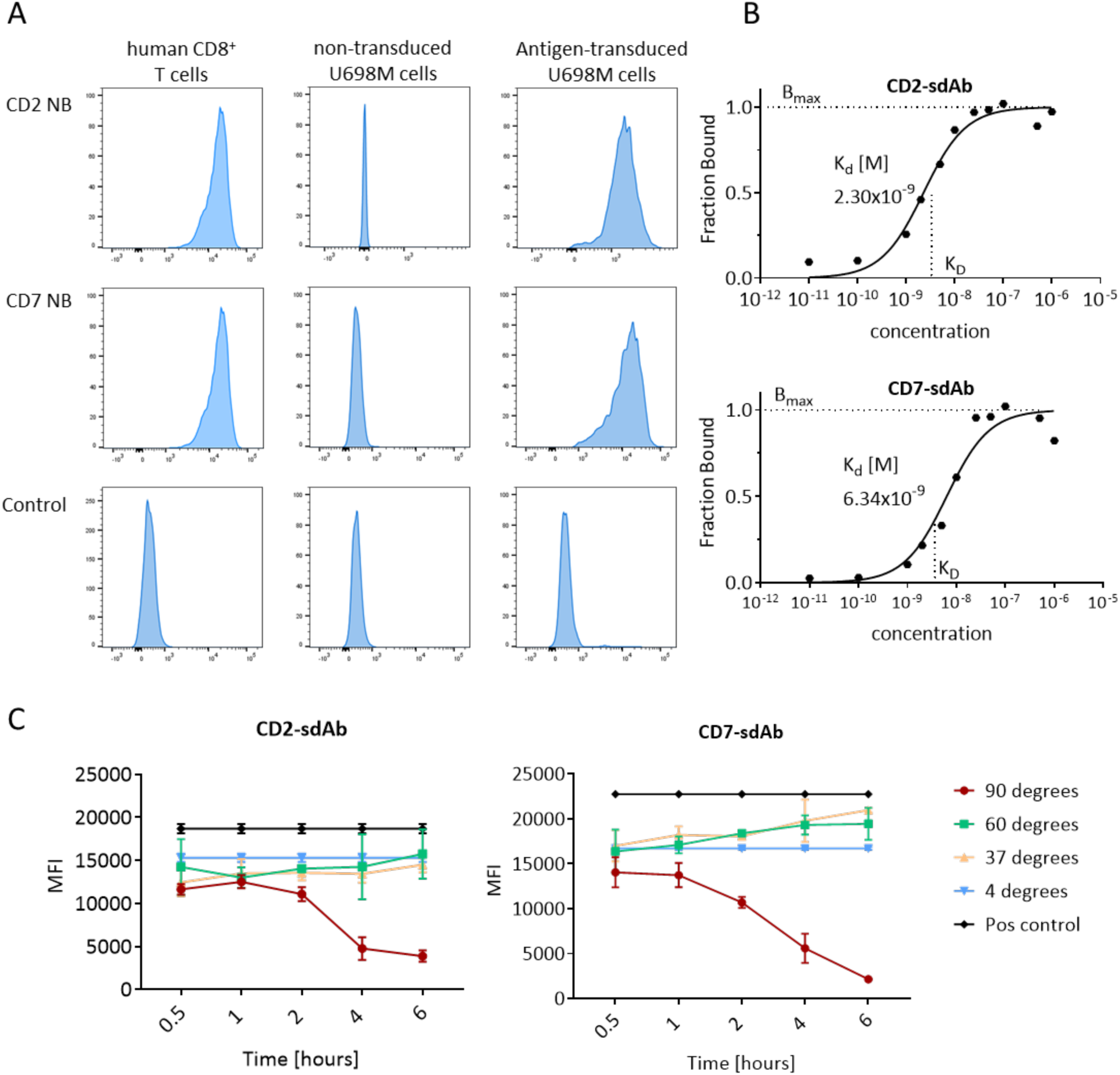
CD2-sdAb and CD7-sdAb show high specificity, affinity and a high degree of thermal binding stability to their respective antigens. **A**, Flow cytometry-based determination of binding of CD2- and CD7-sdAb. Binding capabilities were assessed on human CD8^+^ T cells (left), on CD2- and CD7-negative wild type U-698-M tumor cells which served as a negative control (middle); and on U-698-M tumor cells retrovirally transduced with CD2 (upper graph) and CD7 (middle graph) respectively (right). **B**, Binding affinities of CD2- and CD7-sdAb on human CD8^+^ T cells. Determined affinity binding curves for CD2-sdAb (down) and CD7-sdAb (up) at different concentration on human CD8^+^ T cells. Binding of CD2-sdAb and CD7-sdAb is shown as a fraction of the maximum specific binding (Bmax), measurements were performed in triplicates and data depicted as semi-logarithmic plots. **C**, Analysis of thermal binding stability for each sdAb after incubation at 4° C, 37° C, 60° C and 90° C for 0.5 h, 1 h, 2 h, 4 h and 6 h respectively. After each time interval, sdAb were used for flow cytometry-based binding analysis on CD2- and CD7-positive Jurkat cells and resulting MFIs are depicted with mean +/- standard deviation (sd).

In addition to the binding specificity, thermal binding stability was analyzed to ensure functional binding capabilities of the constructs during high temperatures that are used for downstream modifications such as radiolabeling. After incubation at 4, 37, 60 and 90° C for up to six hours, sdAb were co-incubated with the highly CD2 and CD7 positive acute T-cell leukemia cell line Jurkat to magnify even small differences in thermostability. Subsequent flow cytometry analysis showed that CD2-sdAb and CD7-sdAb could maintain stable antigen binding for up to six hours at 60° C. Only after two hours at 90° C for CD7-sdAb and four hours for CD2-sdAb did we observe a loss of thermal stability in the form of reduced antigen-binding ability (Fig. 2C).

### CD2-sdAb and CD7-sdAb act as inert surrogate markers with no effect on T-cell functions as cytotoxicity and cytokine production in vitro

Previous work by Mayer & Mall et al. [39] has highlighted the importance of thorough and detailed analysis regarding the effect of immuno-tracers on the functionality of their respective target cells. To make sure that CD2-sdAb and CD7-sdAb function as inert T-cell tracers and do not affect T-cell functionality, we analyzed the T-cell cytokine production profile after sdAb binding as well as dynamic T-cell anti-tumor efficacy.

Co-incubation of T cells with CD2-sdAb and CD7-sdAb did not affect cytokine levels compared to the control R3b23-sdAb across three different tumor cell lines transduced with HLA-B*07:02 (ML2-B7, NB4-B7, HL60-B7). To comprehensively assess potential impacts on T-cell functionality, we simultaneously analyzed the cytokine profiles of IFNγ, IL2, GM-CSF, and TNFα for each sdAb. Across all tested conditions, no significant alterations in cytokine secretion were observed, indicating that neither CD2-sdAb nor CD7-sdAb interfered with T-cell activation or effector function (Fig. 3A). These findings further reinforce the safety of CD2- and CD7-targeting sdAbs, demonstrating that they do not induce unintended cytokine-mediated cell signaling upon binding in vitro.

**Fig. 3.**
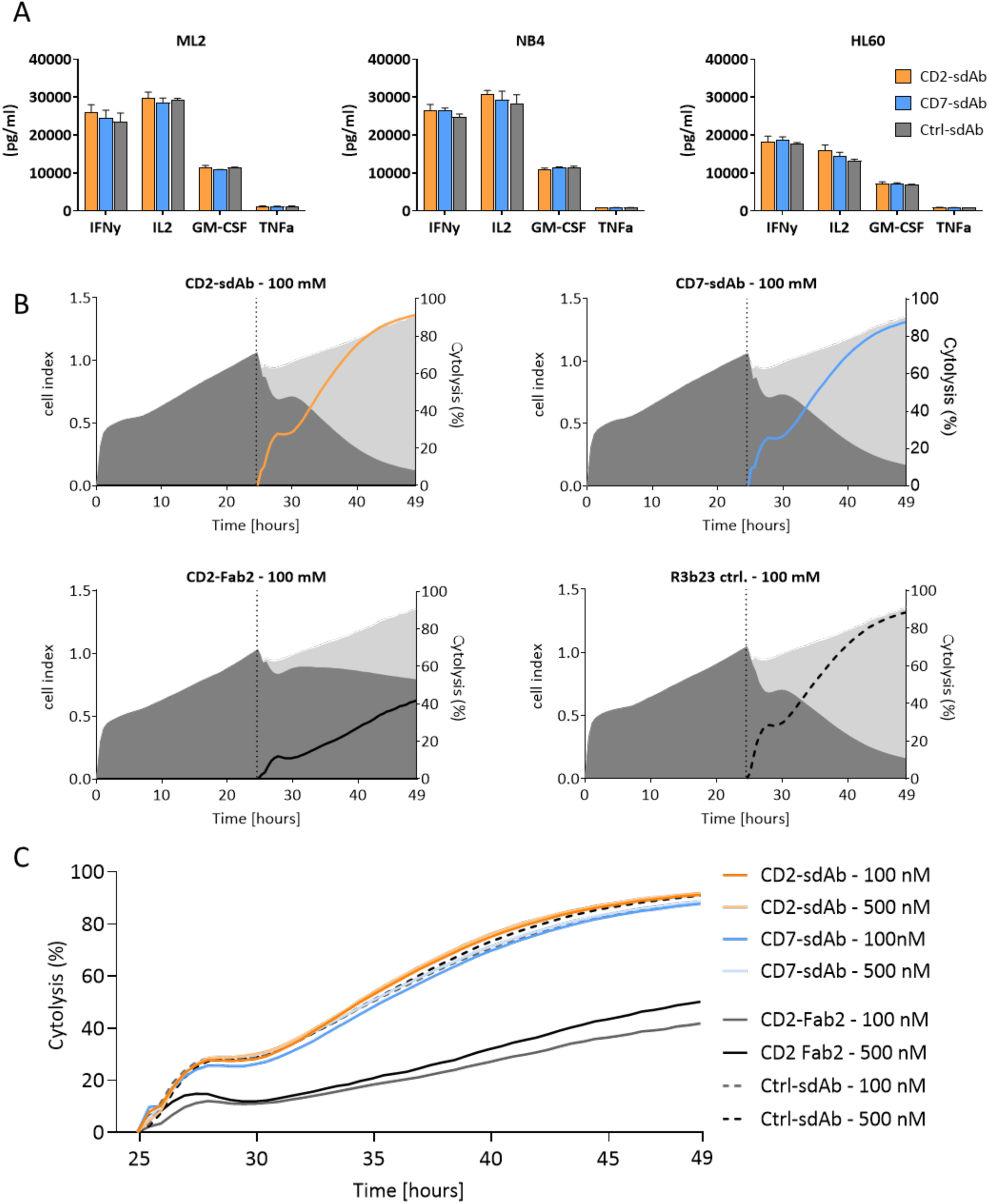
Co-incubation of CD2-sdAb and CD7-sdAb exhibit no impairment of cytotoxicity of T cells and do not alter their cytokine secretion profile. **A**, ELISA-based analysis of IFNγ, IL-2, GM-CSF and TNFα-secretion of TCR-transduced CD8^+^ T cells after 24 h incubation with either CD2-sdAb (orange), CD27-sdA (blue) and R3b23-sdAb as a control (grey) in three different tumor cell lines that have been transduced with HLA-B*07:02 (ML2-B7, NB4-B7, HL60-B7). Results of triplicates are depicted with mean +/- standard deviation (sd). **B**, Proliferation of target tumor cells (624.38 Mel) was monitored by impedance measurement every 30 min for 24 h, after which TCR-transgenic CD8^+^ T cells and the respective antibody-derived constructs were added to the co-culture. 624.38 Mel cells were seeded at a concentration of 75,000 cells / well and after 24 h 75,000 CD8^+^ T cells / well were added together with 100 nM of CD2-sdAb (top left), CD7-sdA (top right), anti-CD2-F(ab’)_2_ (bottom left) and anti-R3b23-sdAb (bottom right). The left y-axis shows cell-index values of tumor cells, indicated by grey planes, while the right y-axis shows the percentage of lysed tumor cells, indicated by colored lines. **C**, Depiction of dynamic target-cell lysis after addition of CD8^+^ T cells and antibody-derived constructs, which have been added at a concentration of 100 nM and 500 nM.

While previously used in vitro assays did not indicate significant impairment of T-cell functionality after CD2-F(ab′)_2_ binding [39], the detrimental effects of CD2 binding on T-cell function only became evident in subsequent in vivo studies. This was because the previously used in vitro assays lacked the sensitivity to detect functional impairments, necessitating in vivo validation to reveal the effects of CD2-F(ab′)_2_ binding. This underscores the need for more sensitive in vitro approaches to assess potential impairments in T-cell activity before progressing to in vivo studies.

To address this, we optimized the xCELLigence assay to dynamically monitor potential functional impairments caused by CD2- and CD7-sdAb binding. This system utilizes cellular impedance to continuously assess the viability of MPO5-positive target tumor cells (624.38 Mel) in the presence of MPO5-specific TCR2.5D6-transduced T cells and the respective sdAbs. The anti-R3b23-sdAb served as a negative control, while the OKT11 (anti-CD2)-F(ab′)_2_ fragment - previously shown to impair T-cell cytotoxicity [39] - was included as a positive control. As expected, TCR-transduced T cells achieved 90% tumor lysis in the presence of the negative control (Fig. 3B, bottom right), whereas co-incubation with the OKT11 (anti-CD2)-F(ab′)_2_ fragment significantly reduced cytotoxic efficacy, resulting in only 40% tumor-cell lysis (Fig. 3A, bottom left).

In contrast, neither CD2-sdAb (Fig. 3B, top left) nor CD7-sdAb (Fig. 3B, top right) impaired T-cell mediated lysis. Even when their concentrations were increased from 100 nM to 500 nM, T-cell cytotoxicity remained unaffected compared to the anti-R3b23-sdAb control (Fig. 3C). Notably, while Mayer et al. required in vivo assays to reveal the impairing effect of CD2-F(ab′)_2_, the xCELLigence assay allowed us to already detect these differences in vitro with high sensitivity. These results highlight under improved detection sensitivity that CD2- and CD7-sdAb binding does not compromise T-cell cytotoxicity in vitro.

Thus, we have demonstrated through comprehensive in vitro analyses, that we do not see evidence of CD7-sdAb and CD2-sdAb to impair T-cell functionality nor impact their cytokine-based cell-signaling upon binding in vitro.

### Cytotoxic efficacy of TCR-transduced T cells in vivo is not affected and impaired by injection of anti-CD2 and anti-CD7 sdAb

To confirm that CD7-sdAb and CD2-sdAb act as inert pan T-cell tracers not just in vitro but also in vivo, we investigated if the application of either construct would impact the ability of TCR-transduced T cells to fully reject xenograft tumors in a previously described myeloid sarcoma model. For that, NSG mice were subcutaneously injected with two tumors, one MHC-matched tumor (ML2-B7) that expressed the MPO-related peptide to be recognized by 2.5D6iRFP CD8^+^ T cells, and one that bears the irrelevant restriction element HLA-B15 and serves as an internal control (ML2-B15). After tumor onset, TCR-transduced T cells were intravenously injected and three days later mice received 5 µg of unconjugated tracer via their tail veins (Fig. 4A). Since previous work by Mall & Mayer et al. [39] has shown that application of anti-CD2-F(ab’)_2_ (clone OKT11) severely impaired T-cell function in vivo, we utilized two different anti-CD2-F(ab’)_2_ clones (OKT11 and RPA-2.10) to serve as positive controls, while groups receiving either PBS or the irrelevant anti-R3b23-sdAb functioned as negative controls. After tumors initially continued to grow for two to three days post T-cell injection, T cells in mice that received either PBS or anti-R3b23-sdAb were able to reject MHC-matched tumors effectively (Fig. 4C), while the internal control-tumors kept growing unaffectedly (Fig. 4B). Contrary to this, administration of both anti-CD2-F(ab’)_2_ clones (OKT11 and RPA-2.10) compromised the functionality of T cells and led to failure of tumor rejection, with treated animals demonstrating continued tumor grow after an initial plateau-phase (Fig. 4B). However, application of CD7-sdAb and CD2-sdAb did not result in loss of T-cell functionality or alter the dynamics of tumor rejection, which were completely rejected by day eleven as compared to mice receiving PBS or anti-R3b23-sdAb (Fig. 4B). The significant difference between the effects of anti-CD2-F(ab’)_2_ and CD2-sdAb on T-cell functionality, in vitro and in vivo, provides a strong indication that in contrast to anti-CD2-F(ab’)_2_ the particular binding capabilities of CD2-sdAb allow for specific binding of CD2 on T cells without alteration of their function. In addition, similar to the previously described anti-CD7-F(ab’)_2_ [39], the CD7-sdAb showed no functional impairment in vivo.

**Fig. 4.**
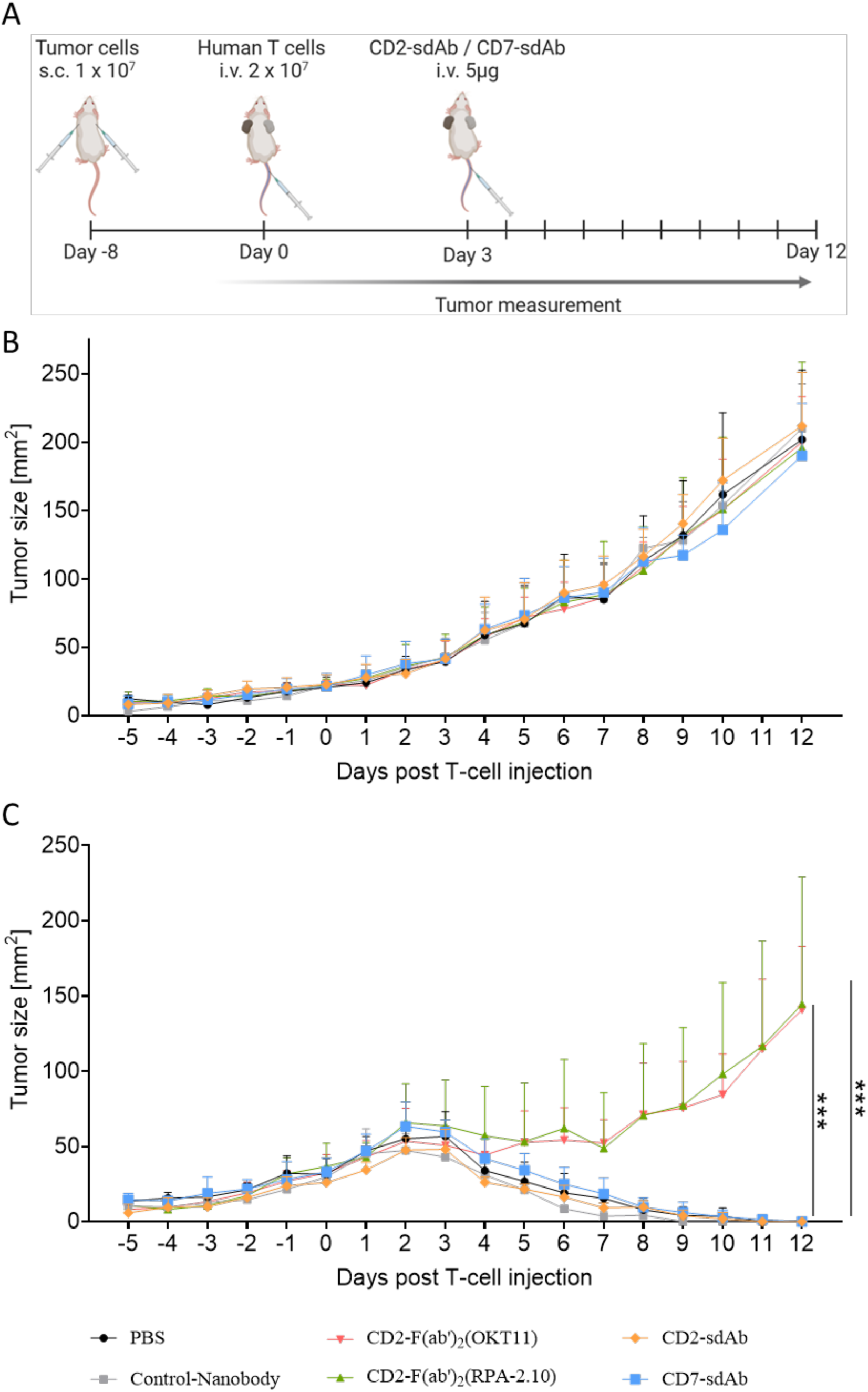
Application of CD2-sdAb and CD7-sdAb did not impair T-cell function in vivo. **A**, Experimental setup of in vivo tumor rejection model for CD8^+^ T cells. NSG mice were subcutaneously (s.c.) injected with ML2-B7 cells in the right flank and ML2-B15 cells in the left flank. After eight days, TCR-transgenic human CD8^+^ T cells were injected intravenously (i.v.) through the tail vein, followed three days later by i.v. injection of R3-b23-sdAb, OKT11, RPA, CD2-sdAb or CD7-sdAb. Tumor growth was monitored from tumor onset until the end of experiment at day twelve. **B, C,** Monitoring of tumor growth kinetics of ML2-B15 (B) and ML2-B7 tumors (C) in NSG mice. On day 0, eight days after subcutaneous tumor injection, mice were intravenously injected with TCR 2.5D6-transgenic CD8^+^ T cells and three days later with either PBS, R3b23-sdAb, CD2-F(ab’)_2_ (OKT11), CD2-F(ab’)_2_ (RPA-2.10), CD2-sdAb or CD7-sdAb. Kinetics of tumor growth were monitored daily for twelve days post T-cell injection. The experiment was ended at day twelve, at which point all relevant non-control tumors had been fully rejected. Tumor sizes are shown in mm^2^ and mean values and SDs are depicted for each group of mice. PBS n = 4, R3b23-sdAb n = 3, CD2-F(ab’)_2_ (OKT11) n = 6, CD2-F(ab’)_2_ (RPA-2.10) n = 4, CD2-sdAb n = 5, CD7-sdAb n = 5. Significance was calculated using Mann-Whitney test (* p ≤ 0.05, ** p ≤ 0.01, *** p ≤ 0.001)

### In vivo imaging using ^68^Ga-NOTA-CD2- and ^68^Ga-NOTA-CD7-sdAb enables tracking of intravenously injected human CD8+ T cells, showing a distinct signal at the HLA-matched tumor site

Since thorough characterization of both sdAb has proven them to be both highly specific while also not affecting their target cells in terms of functionality, both constructs were used to determine their aptitude for in vivo T-cell imaging.

To assess the ability of ⁶⁸Ga-NOTA-CD2- and ⁶⁸Ga-NOTA-CD7-sdAb to track human T cells in vivo, a tumor-bearing NSG mouse model was established (Fig. 5A). On day -8, ML2-B7 tumor cells, which express the HLA-B07:02 molecule and are recognized by the infused T cells, were subcutaneously injected into the right shoulder of mice (1 × 10⁷ cells per mouse). To serve as an internal control for non-specific T-cell accumulation, a control tumor line, ML2-B15, which lacks HLA-B07:02, was injected into the left shoulder of the same mice. On day 0, human CD8⁺ T cells transduced with an HLA-B*07:02-restricted TCR were intravenously injected (2 × 10⁷ cells per mouse). Five days later, mice were administered ⁶⁸Ga-NOTA-CD2-sdAb or ⁶⁸Ga-NOTA-CD7-sdAb via intravenous injection. As an additional control, a separate cohort of mice received ⁶⁸Ga-NOTA-R3b23-sdAb, a non-T-cell-targeting sdAb, to evaluate non-specific accumulation. One hour post-injection (p.i.), mice underwent PET/MR imaging to assess tracer distribution and T-cell localization (Fig. 5A).

**Fig. 5.**
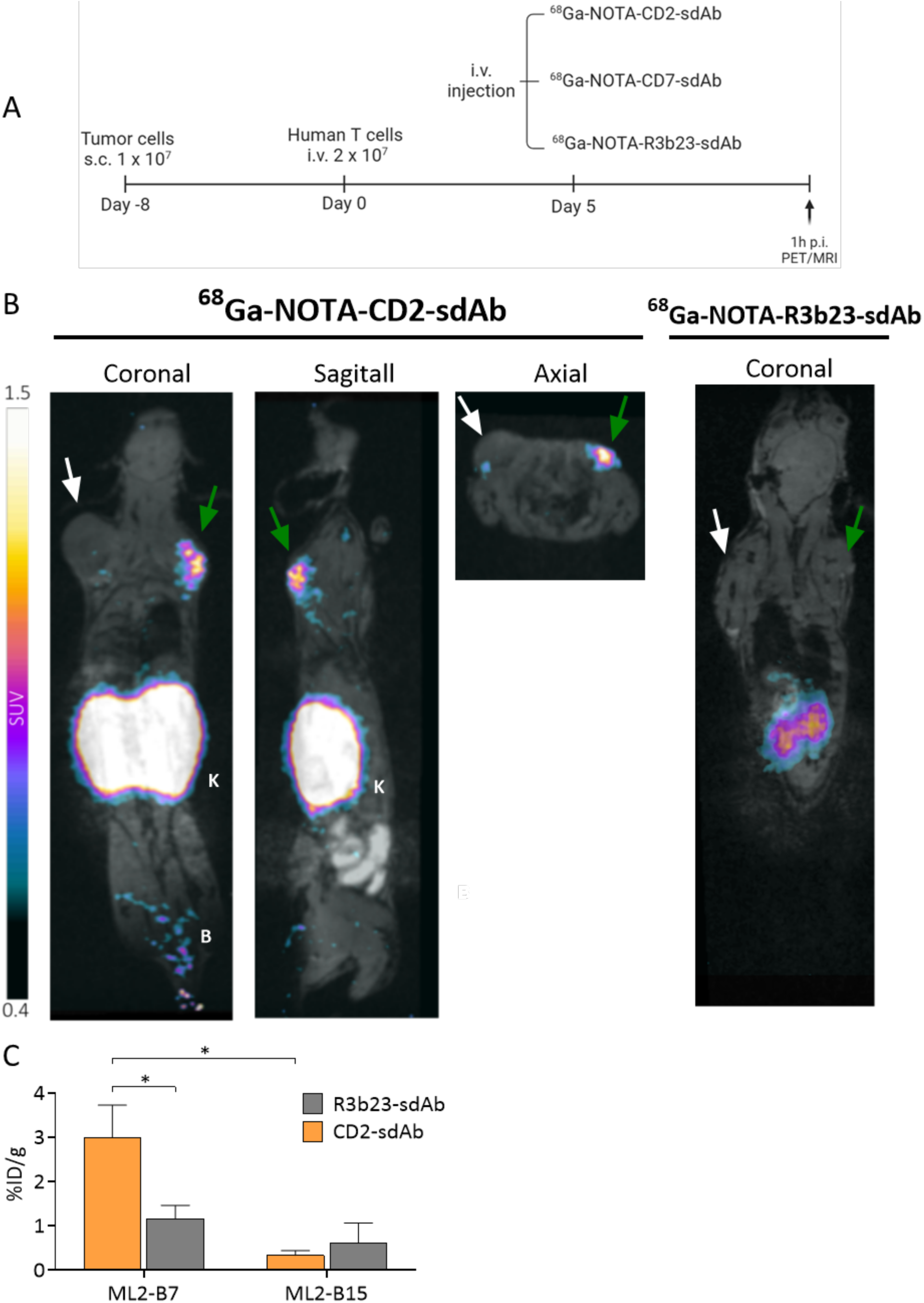
In vivo tracking intravenously injected TCR-transgenic CD8+ T cells in ML2-B7 tumors using ^68^Ga-NOTA-CD2-sdAb. **A**, Experimental setup for in vivo imaging of ⁶⁸Ga-NOTA-labeled sdAb in NSG mice. ML2-B7 (HLA-matched) and ML2-B15 (irrelevant control) tumors were subcutaneously injected, followed by intravenous injection of TCR-transgenic CD8⁺ T cells (2 × 10⁷ per mouse). Three groups of five mice received either ⁶⁸Ga-NOTA-CD2-sdAb, ⁶⁸Ga-NOTA-CD7-sdAb, or ⁶⁸Ga-NOTA-R3b23-sdAb, and PET/MRI scans were performed one hour post-injection (p.i.). **B**, PET/MRI images acquired one h p.i. of mice injected i.v. with ^68^Ga-NOTA-CD2-sdAb (left) or ^68^Ga-NOTA-R3b23-sdAb (right). Exemplary mice are shown in coronal, sagittal and axial orientation (A) or just in coronal orientation (B). 2 x 10^7^ TCR-transgenic CD8^+^ T cells were injected i.v. and the injected dose of applied tracer was 13 ±1MBq per mouse. Scale bar is represented as standardized uptake value (SUV), 0.4 – 1.5 SUV. B = Bladder, K = Kidney. Green arrow = ML2-B7 tumor, white arrow = ML2-B15 tumor. **C**, Biodistribution of ^68^Ga-activity in ML2-B7 and -B15 tumors for mice receiving CD2-sdAb or R3b23-sdAb 1.5 h post injection and after previous PET/MRI image acquisition. Mean %ID/g ± SD is depicted for each group of mice. R3b23-sdAb n = 4, CD2-sdAb n = 4. Significance was calculated using Mann-Whitney test (* p ≤ 0.05).

PET/MR imaging demonstrated a clear accumulation of ⁶⁸Ga-NOTA-CD2-sdAb at the ML2-B7 tumor site, whereas no notable signal was observed at the ML2-B15 control tumor (Fig. 5B). Tracer uptake in the ML2-B7 tumor was consistently visible across coronal, sagittal, and axial imaging planes, with additional uptake primarily restricted to the kidneys, reflecting renal clearance of the tracer. Mice injected with the control ⁶⁸Ga-NOTA-R3b23-sdAb displayed signal accumulation exclusively in the kidneys, with no detectable uptake in either tumor, confirming the specificity of CD2-targeted imaging (Fig. 5B).

Ex vivo biodistribution analysis further corroborated these findings. The injected dose per gram of tumor tissue (%ID/g) was significantly higher in ML2-B7 tumors of mice receiving ⁶⁸Ga-NOTA-CD2-sdAb (mean 3.0 %ID/g) compared to those receiving ⁶⁸Ga-NOTA-R3b23-sdAb (mean 1.2 %ID/g) (Fig. 5C). Additionally, in mice injected with ⁶⁸Ga-NOTA-CD2-sdAb, tracer uptake in ML2-B7 tumors (3.0 %ID/g) was significantly higher than in ML2-B15 tumors (0.3 %ID/g), further highlighting the specificity of the tracer for HLA-matched tumor sites. Together, these findings indicate that ⁶⁸Ga-NOTA-CD2-sdAb enables effective in vivo tracking of intravenously injected human CD8⁺ T cells, showing specific localization at the HLA-matched tumor site.

Similarly, in vivo evaluation of ⁶⁸Ga-NOTA-CD7-sdAb for tracking intravenously injected CD8⁺ T cells revealed results comparable to those observed with ⁶⁸Ga-NOTA-CD2-sdAb, albeit with slightly lower overall signal intensity. PET/MR imaging demonstrated strong and specific accumulation of ⁶⁸Ga-NOTA-CD7-sdAb at the ML2-B7 tumor site, whereas no notable signal was detected at the ML2-B15 control tumor (Fig. 6A). The specific tracer uptake in ML2-B7 tumors was consistently visible across coronal, sagittal, and axial orientations throughout the imaging process.

**Fig. 6.**
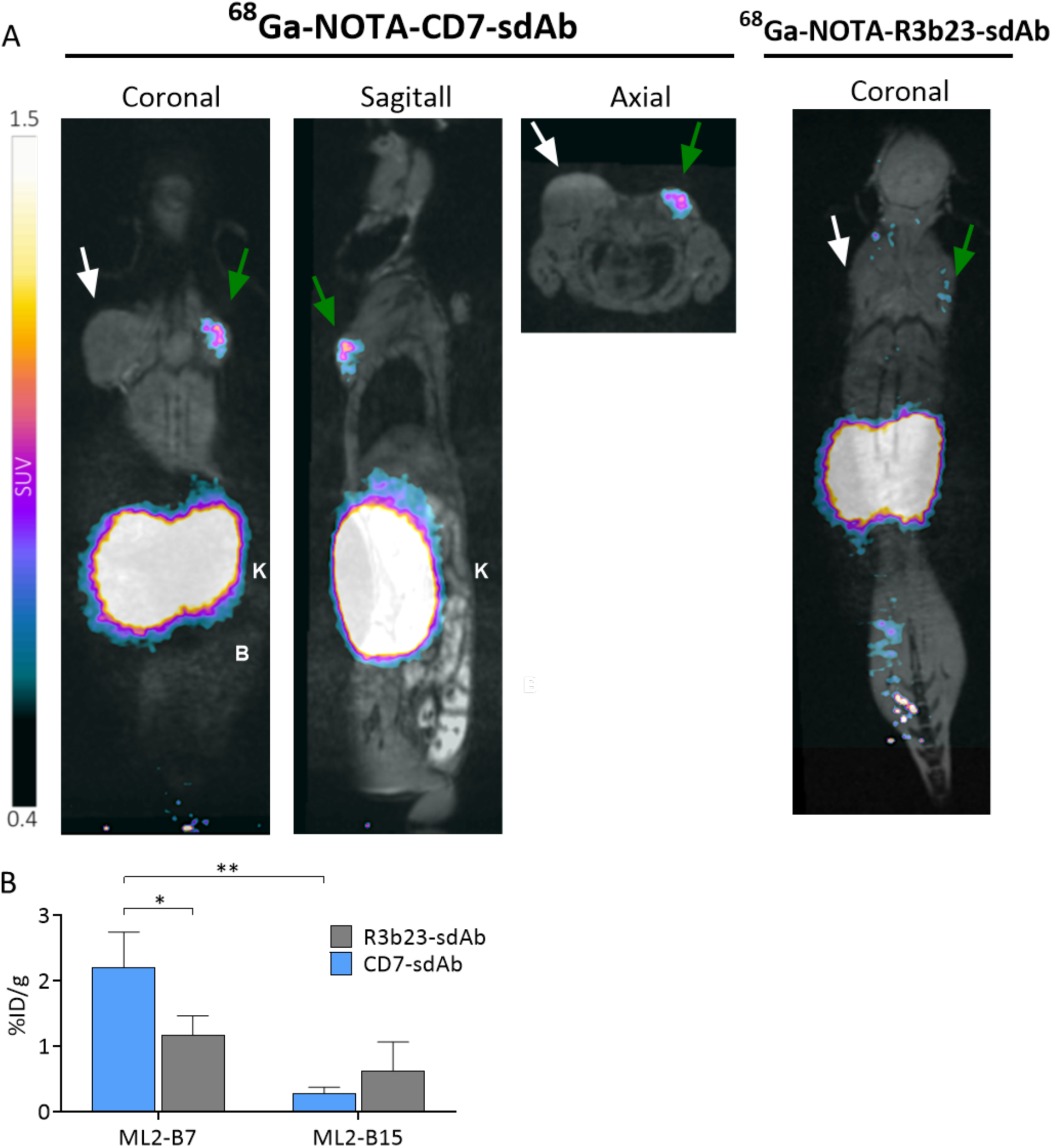
In vivo tracking intravenously injected TCR-transgenic CD8+ T cells in ML2-B7 tumors using ^68^Ga-NOTA-CD7-sdAb. **A**, PET/MRI images acquired one h p.i. of mice injected i.v. with ^68^Ga-NOTA-CD7-sdAb (left) or ^68^Ga-NOTA-R3b23-sdAb (right). Exemplary mice are shown in coronal, sagittal and axial orientation (left) or just in coronal orientation (right). 2 x 10^7^ TCR-transgenic CD8^+^ T cells were injected i.v. and the injected dose of applied tracer was 14 ±1MBq per mouse. Scale bar is represented as standardized uptake value (SUV), 0.4 – 1.5 SUV. B = Bladder, K = Kidney. Green arrow = ML2-B7 tumor, white arrow = ML2-B15 tumor. **B**, Biodistribution of ^68^Ga-activity in ML2-B7 and -B15 tumors for mice receiving CD7-sdAb or R3b23-sdAb 1.5 h post injection and after previous PET/MRI image acquisition. Mean %ID/g ± SD is depicted for each group of mice. R3b23-sdAb n = 4, CD7-sdAb n = 5. Significance was calculated using Mann-Whitney test (* p ≤ 0.05, ** p ≤ 0.01).

Ex vivo biodistribution analysis further confirmed the PET/MRI data, showing significantly higher tracer accumulation in ML2-B7 tumors of mice injected with ⁶⁸Ga-NOTA-CD7-sdAb compared to ML2-B15 tumors (Fig. 6B). Quantification of tumor-associated activity revealed a mean uptake of 2.2 %ID/g in ML2-B7 tumors, whereas the ML2-B15 tumors exhibited only 0.3 %ID/g, demonstrating a clear and statistically significant preference for the HLA-matched tumor environment.

To further assess the systemic distribution of the tracer, biodistribution analyses of various organs were performed following PET/MR imaging. As expected, the highest tracer uptake was observed in the kidneys, reflecting renal clearance, with minor uptake in the bladder and heterogeneous accumulation in the liver (Figure 7). These findings underscore the specificity of ⁶⁸Ga-NOTA-CD7-sdAb for detecting human CD8⁺ T cells at the tumor site and reinforce its potential as a valuable imaging tool for monitoring T-cell localization in vivo.

**Figure 7:**
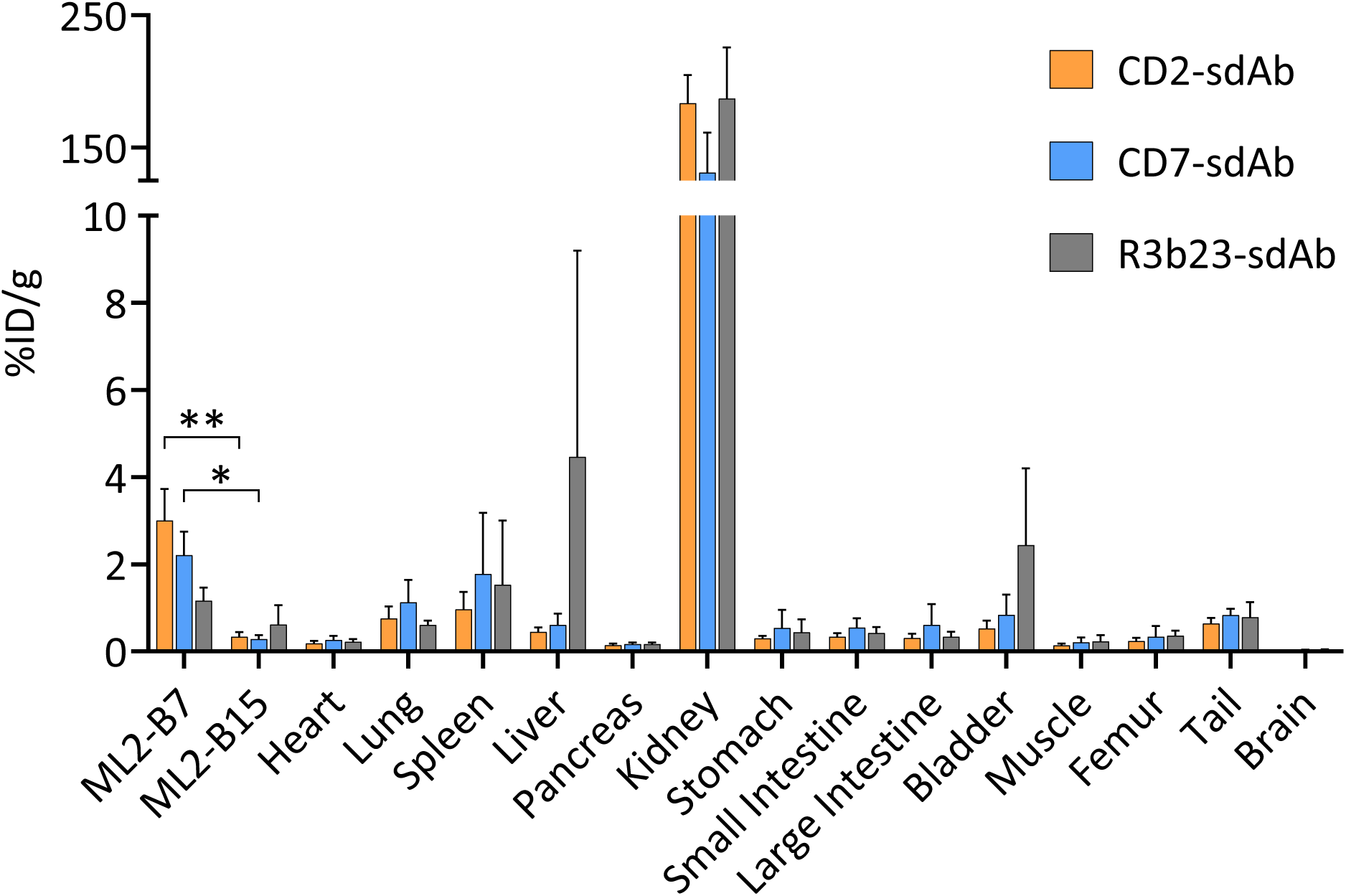
Ex vivo biodistribution of ^68^Ga-radiolabeled CD2- and CD7-sdAb. Biodistribution of ^68^Ga-activity in various organs from mice 1.5 h post injection of the respective sdAb and following previously described PET/MRI analysis. Mean %ID/g ± SD is depicted for each group of mice. R3b23-sdAb n = 4, CD2-sdAb n = 4, CD7-sdAb n = 5. Significance was calculated using Mann-Whitney test (* p ≤ 0.05, ** p ≤ 0.01).

## DISCUSSION

### Evaluation of CD2 and CD7 as Target Antigens for ImmunoPET Imaging

Establishing a preclinical T-cell tracer with strong potential for clinical translation requires selecting a suitable target antigen, a well-characterized probe, and an appropriate radionuclide [49]. Since the choice of radionuclide for CD2- and CD7-sdAb tracers was determined by our collaborators in the Department of Nuclear Medicine, this discussion focuses on target antigen selection and probe characterization.

Over the past decade, CD8 has been the primary focus for T-cell-based immunoPET imaging, with multiple preclinical studies demonstrating its utility in visualizing CD8^+^ T cells in response to immunotherapy [50]. The only clinically tested T-cell tracer to date, [^89^Zr]Zr-IAB22M2C, specifically targets CD8^+^ T cells and has shown safety and feasibility in a first-in-human trial [51]. However, CD8-based tracers are limited to monitoring CD8^+^ T cells, whereas immunotherapy responses can also involve CD4^+^ T cells [5, 52]. Additionally, novel therapies increasingly leverage the distinct roles of CD4^+^ T cells, such as CD4^+^ CAR T cells, which demonstrated enhanced long-term tumor eradication compared to CD8^+^ CAR T cells in a preclinical leukemia model [38]. The FDA-approved CAR T-cell therapy lisocabtagene maraleucel also includes both CD4^+^ and CD8^+^ T-cell components, underscoring the importance of comprehensive immune monitoring [36]. To enable a more encompassing assessment of T-cell responses, we evaluated the pan-T-cell markers CD2 and CD7 as alternative immunoPET targets.

CD2 was first identified as a pan-T-cell antigen by Sanchez-Madrid et al. [53] and has been widely used in flow cytometry [54] and immunohistochemistry [55]. Beyond its role as a T-cell marker, CD2 functions as an important adhesion and costimulatory molecule during T-cell activation [56], mediating interactions with antigen-presenting cells through binding to its ligand CD58 (LFA-3) [57]. This interaction contributes to immunological synapse formation and enhances T-cell signaling [58], making CD2 a relevant and dynamic marker of immune activation. A key advantage of CD2 is its elevated expression on activated T cells, as repeatedly reported [59, 60], including our own findings [39]. This feature makes CD2 an attractive target, as it allows for visualization of both the overall T-cell population and specifically activated T cells.

Although CD2 is predominantly expressed on T cells, it is also found on NK cells [61] and subsets of dendritic cells [62], which could contribute to off-target tracer binding. However, imaging signals in these cell populations may be also of interest [63] although their quantity may be below the detection limit. Thus, while non-T-cell binding is a consideration, CD2 expression on NK and dendritic cells may still provide clinically relevant information [64]. Future studies, such as humanized mouse models, may be helpful to determine the impact of these additional cell types on imaging specificity.

CD7 shares similarities with CD2 in that it is expressed on T cells, thymocytes (Barcena et al., 1993), and NK cells [65]. Functionally, CD7 is a transmembrane glycoprotein belonging to the immunoglobulin superfamily and plays a role in T-cell interactions and early lymphocyte signaling [66]. It is involved in the regulation of T-cell activation and cytokine production [67], although its precise function remains less well defined than that of other costimulatory molecules. CD7 differs from CD2 in its higher expression on naïve and memory T cells and for showing only a modest increase in expression upon T-cell activation [39]. Additionally, while CD2 is absent on early T-cell precursor (ETP) and pro-T cells [68], CD7 is expressed throughout all stages of T-cell development, which could result in increased tracer accumulation in the thymus.

Taken together, while both CD2 and CD7 are viable T-cell targets, CD2’s stronger upregulation on activated T cells and lower expression on early thymocytes make it the preferred antigen for non-invasive T-cell monitoring in immunotherapy settings.

### Specific Binding Site of CD2-sdAb and Implications

Our previous study using a CD2-targeted F(ab′)₂ fragment labeled with [^89^Zr]Zr successfully tracked tumor-specific T cells in a xenograft model but led to severe T-cell depletion and tumor rejection failure [39]. This F(ab′)₂ fragment was derived from the OKT11 monoclonal antibody, which binds the T11.1 epitope of CD2, a constant adhesion domain required for CD2–CD58 interaction [69]. Most commercially available anti-CD2 antibodies also target T11.1 or T11.2 due to their continuous expression, making them useful for therapeutic applications in autoimmune diseases and transplantation [70].

In contrast, CD2-sdAb did not impair T-cell function in vitro or in vivo. Single-domain antibodies (sdAbs) can bind otherwise inaccessible protein clefts due to their unique CDR3 region and hypervariable loops [71, 72]. Thus, CD2-sdAb may bind T11.1 or T11.2 in a non-disruptive manner, preserving T-cell functionality. Alternatively, it may target the primarily activation-induced T11.3 epitope [73], enabling increased specific visualization of activated T cells.

To distinguish between these possibilities, future studies will be needed to precisely identify the binding site and provide a clearer understanding of how target recognition occurs at the molecular level.

### Impact of CD2- and CD7-sdAb Binding on T-Cell Functionality

Assessing potential immunomodulatory effects of tracers is crucial for immunoPET imaging, as mAbs and their derivatives can impair T-cell function and affect therapeutic efficacy potential [74]. Previous work from our group demonstrated that in vivo application of a CD2-targeted F(ab′)_2_ fragment led to severe T-cell depletion and tumor rejection failure [39]. However, in vitro studies failed to properly show and demonstrate a reduction in T-cell cytotoxicity when CD2-F(ab′)_2_ was added to T-cell and tumor co-cultures, highlighting the limitations of static assays.

To enhance preclinical assessment, we transitioned from flow cytometry-based analysis to real-time monitoring via the XCELLigence system. This approach enables continuous evaluation of tumor cell viability by measuring cellular impedance, providing a more sensitive and dynamic assessment of T-cell functionality. Using two CD2-targeted F(ab′)_2_ fragments (OKT11, RPA-2.10) as positive controls, we confirmed significant impairment of T-cell mediated tumor cell killing, consistent with previous in vivo findings (Fig. 3). Subsequent in vivo experiments further validated that while CD2-F(ab′)_2_ constructs led to failed tumor rejection, CD2- and CD7-sdAb had no impact on T-cell functionality (Fig. 4). Based on these results, the xCELLigence assay will be integrated into the workflow to screen potential imaging constructs for T-cell impairing effects before in vivo studies.

Beyond direct functional impairment, targeting T cells with antibodies can also induce cytokine release syndrome (CRS), a severe inflammatory condition characterized by elevated levels of IFNγ, TNFα, IL-6, and other cytokines [75]. In adoptive T-cell therapy, CRS progresses in two phases: initially, T cells release IFNγ and TNFα, which then activate monocytes and macrophages to further amplify cytokine secretion [76–78]. Clinical manifestations include fever, systemic inflammation, and, if untreated, organ failure [79, 80]. While preclinical models cannot fully predict clinical CRS outcomes [81], they are useful for assessing immediate cytokine changes.

To minimize patient risk, we analyzed cytokine profiles (IFNγ, GM-CSF, IL-2, TNFα) in the presence of CD2- and CD7-sdAb across three cell lines. After 24 hours, no significant alterations in cytokine secretion were observed compared to the control R3b23-sdAb (Fig. 3), suggesting that CD2- and CD7-sdAb do not induce unwanted cytokine responses.

### Potential non-specific off-target effects

Selecting both an appropriate target antigen and an optimal tracer construct is crucial for effective immunoPET imaging. Single-domain antibodies (sdAbs) offer several advantages, including low immunogenicity [82], high affinity [83], specificity [84], fast renal clearance [85], and efficient tissue penetration [86]. These properties make sdAbs ideal for rapid target visualization and minimizing patient radiation exposure.

However, due to their small size, sdAbs exhibit high renal and bladder uptake [87], driven by rapid renal clearance [88]. While this ensures a low background signal and quick imaging, it also increases kidney radiation burden [89]. Strategies to reduce renal retention include modifying sdAb C-terminal residues, as removal of a Myc-His-Tag significantly decreased kidney accumulation [90], and blocking megalin-mediated reabsorption using arginine-lysine, monosodium glutamate, or gelofusine solution [91, 92].

Another limitation of sdAbs is rapid blood clearance, potentially reducing target-site binding. To counter this, sdAbs can be modified with polyethylene glycol (PEG) to extend circulation time [93]. A study using a ^89^Zr-labeled anti-CD8 sdAb demonstrated that larger PEG modifications (5–20 kDa) correlated with reduced kidney uptake [94]. However, increasing sdAb size may compromise tissue penetration, a key advantage over larger tracers.

For example, an ^89^Zr-labeled anti-CD8 minibody (~80 kDa) visualized CD8^+^ lesions in a clinical trial but required over 24 hours for sufficient target binding, likely due to its larger molecular weight [51]. In contrast, sdAbs enable rapid target binding and fast clearance, making them ideal for visualizing T cells in immunotherapy settings.

### Suitability of ^68^Ga for ImmunoPET Imaging (or similar)

The selection of radioisotopes in Nuclear Medicine imaging modalities should be based on several characteristics like their half-life, decay modality and energy, as well as their availability. In particular, the physical half-life of the radionuclide must match the expected biological half-life of the vector molecules in vivo, to allow for the optimal imaging results.

Gallium-68 with a half-life of 68 min, decays to stable zinc-68, 89% through positron emission with a mean energy of 836 keV (E^β+^,max=1.89 MeV) and is then perfectly suitable for the radiolabelling of small biomolecules such as single domain antibodies. In contrast to common PET isotopes, ^68^Ga is normally not produced by on-site cyclotrons but is obtained from ^68^Ge/^68^Ga radioisotope generators (t_1/2_(^68^Ge) = 270.8 days). Due to its low radiation energy, low or no radiotoxicity should be expected on immune cells, for example within T cells tracking applications, but also on other tissue and cells. Its incorporation in biological constructs occurs by a complexation reaction at room or mild temperatures via chelators previously conjugated to the proteins. The soft labelling conditions complete its suitability with single domain antibodies in ImmunoPET imaging modalities. Zhao et al. have reported the use of ^68^Ga used with a single domain antibody based tracer to track human CD8^+^ T-cells in vivo via ImmunoPET [95]. The radiotracer [^68^Ga]Ga-NOTA-SNA006a, designed to target the human CD8 antigen, was successfully synthesized with high radiochemical purity and strong affinity. It demonstrated promising performance in tracking human CD8+ T cells in mouse models, outperforming other candidates.

### CD2- and CD7-sdAb as ImmunoPET Imaging Tracers

After CD2- and CD7-sdAb were thoroughly characterized in terms of target-antigen specificity, binding affinity, and thermal stability, and it was confirmed that their binding did not impair T-cell functionality in vitro or in vivo, both tracers were evaluated for their suitability in PET/MR imaging.

For this, NSG mice were subcutaneously injected with ML2-B7 and ML2-B15 tumor cells to mimic a responder (ML2-B7) and non-responder (ML2-B15) tumor. After tumor engraftment, 2 × 10^7^ TCR-transgenic CD8^+^ T cells were intravenously injected, followed by administration of ^68^Ga-NOTA-CD2-sdAb or ^68^Ga-NOTA-CD7-sdAb via the tail vein. PET/MR imaging performed one-hour post-injection demonstrated distinct tracer accumulation at the ML2-B7 tumor site, with the expected physiological uptake in the kidneys (Fig 5B, 6A). No detectable signal was observed at the irrelevant ML2-B15 control tumor in each experiment, confirming target specificity.

In sum, all mice showed clear tracer accumulation at ML2-B7 tumors, while no uptake was observed at ML2-B15 tumors or in non-target tissues apart from the kidneys and bladder. Biodistribution analysis confirmed these imaging results, revealing significantly higher tracer uptake in ML2-B7 tumors compared to ML2-B15 tumors and R3b2-sdAb controls for both ^68^Ga-NOTA-CD2-sdAb and ^68^Ga-NOTA-CD7-sdAb. These findings demonstrate that both CD2- and CD7-targeting tracers effectively visualize adoptively transferred T cells in vivo, highlighting their potential for non-invasive T-cell tracking in immunotherapy settings.

### Conclusion and outlook

We developed two novel sdAb-based tracers, CD2-sdAb and CD7-sdAb, for immunoPET imaging of T cells in immunotherapy. Targeting the pan T-cell markers CD2 and CD7 enables visualization of the full spectrum of T-cell responses, overcoming limitations of monitoring restricted T-cell subsets. Extensive validation confirmed target specificity, ensuring high tracer accumulation while minimizing off-target binding for improved patient safety.

To assess potential T-cell impairment after tracer binding, we implemented the xCELLigence assay alongside cytokine profiling and in vivo tumor rejection studies. None of these analyses showed T-cell dysfunction, supporting the safety of both tracers. Beyond evaluating purity, binding affinity, specificity, and thermostability, we tested their ability to track adoptively transferred human T cells in a xenograft mouse model. PET/MRI scans one-hour post-injection of ^68^Ga-labeled CD2- and CD7-sdAb demonstrated highly specific tumor-site accumulation without unexpected off-target signals.

These pan T-cell tracers successfully differentiated between responding and non-responding tumors while preserving T-cell function, making them promising surrogate markers and therefore tools for guiding immunotherapeutic decisions. To advance toward clinical translation, further safety evaluations are needed, including pharmacokinetics and toxicology studies. Since traditional rodent models are unsuitable for assessing human-specific tracers, humanized mouse models may provide a viable approach for toxicity assessments, ensuring a robust preclinical foundation for regulatory approval.

Beyond diagnostic applications, the CD2- and CD7-sdAb tracers hold strong theranostic potential for T-cell lymphomas, where their diagnostic use in immunoPET could be paired with therapeutic radionuclide labeling for targeted treatment. This theranostic strategy would allow patient selection, response monitoring, and therapy delivery to be combined within the same sdAb platform. Ultimately, it may may further be extended through alternative payloads or engineered bispecific sdAbs to broaden platform applications.

## DECLARATIONS

### Acknowledgments

This work was supported by a grant from the Deutsche Forschungsgemeinschaft (SFB824/C10) (A.M.K., C.D.) and TRR338/A03 (A.M.K.). The authors thank, Sybille Reeder, Markus Mittelhäuser, Hannes Rolbieski for excellent other technical support and Franz Hagn for support conducting the MS analyses.

### Authors’ contributions

D.G., and A.M.K. designed studies; D.G., T.K. and S.B. performed in vitro and in vivo experiments; L.R. performed radiolabeling, HPLC analysis and supported in vivo studies; D.G., L.R., T.K., S.B., C.D. and A.M.K analyzed and interpreted experimental data; D.G., L.R. and A.M.K. created the figures; D.G., L.R., C.D. and A.M.K. wrote, reviewed and/or revised the manuscript; W.W., F.B., C.D. and A.M.K. supported with administrative, database construction, technical and material, C.D. and A.M.K. supervised the study.

